# Manipulation-based skills for anthropomorphic human-arm system based on integrated ANFIS and vector calculus

**DOI:** 10.1101/2020.02.10.941344

**Authors:** LE Than, LE An

## Abstract

In this paper, we focus on developing our new approaches to solve Inverse Kinematics using Vector Calculus and integrating the ANFIS module. Specifically, these approaches are evaluated in term of accuracy and performance on the 5-DOF robotic arm model inspired by the human arm structure. Evaluation for our new approaches are described detail in this paper and therefore shows the efficiency and robustness of our methods and potentially leads a new research direction in modeling kinematics of anthro-pomorphic robotic arm.

## 1. Introduction

Nowadays, mobile robot manipulators and humanoid robot are one of the most active research areas in robotics community. The goal of these areas is to develop robust manipulation and dexterous grasp model to deal with the high-dimension task-space scenarios as well as safety behaviors in human-robot interaction. Recently, to tackle with the mentioned goal, many bio-inspired robotics arm models are proposed. For instance, a human friendly robotic arm with hybrid actuators is proposed in [3] to guarantee safe behavior in human-robot interaction; a simulation system to analyze grasping of human hand model is presented in [4], DLR’s anthropomorphic hand arm [5] was developed with new concepts on variable stiffness joints in order to protect the arm in hard-to-predict environments.

After finishing the description of new arm model, the forward and inverse kinematics of the arm model must be developed for control basis. Forward kinematic is relatively easy to solve due to its straight-forward frame transformations. However, Inverse kinematics problem is more challenging, it can have several solutions are possible or no solution in certain circumstances. There are two approaches to solve the problem. First, the numerical approach produces approximately solutions for defined coordinates in Cartesian space, which does not use any pre-trained dataset. However, these methods usually require complex and expensive computation to find the answers. For example, a comprehensive novel of pseudo-inverse Jacobian method is studied in [6] with best computing time 1.672s. Comparing with our proposed method using Vector Calculus, our method outperforms Jacobian method with average 30ms computing time and below 2% error with constraint limitation.

Second solution, recently machine learning approach has shown efficient methods to solve inverse inverse kinematics for robotics arm system with high accuracy using Adaptive Neuro-Fuzzy Inference System (ANFIS) [7], [8]. However, in order to achieve low error, a large training set that covers defined Cartesian space has to be trained. An modified framework that is applied for 4 degree of freedom (DOF) arm is proposed in this document.

In this paper, section II will describe the description of our human arm model. Section III will cover forward kinematics based on Denavit-Hartenberg (DH) parameters and link transformations of the arm. The next section present special description as a basis for vector calculus approach to solve the inverse kinematics and Vector Calculus solution. Then, section V provides the basis of ANFIS and its adaptation for our arm model as a machine learning technique. After that, we will demonstrate the evaluation of both approaches on MATLAB environment to present their reliability and accuracy in solving inverse kinematics problems. Final section is the conclusion extracted from our work and future works.

## 2 Model Description

Kinematics is the science of motion that study motion regardless of the forces that are applied to it. In this paper, we care only about the position and orientation of the linkages. In order to deal with the topology of such a complex manipulator as a humanoid arm, we need to assign frames to its parts. After that, we describe the relationship between them in term of equations.

### 2.1 Spatial Description

The idea is to mimic a sphere joint of human arms by using two revolute joints, whose rotary axes are perpendicular to each other. Each pair of joints acts as a human’s natural joint. Therefore, for simplicity, the arm model consists of five revolute joints, the first two joints corresponds to human shoulder, the second is the elbow and final joint replicates the wrist. This model is suitable for inexpensive devices such as cheap servo motors and can act as a testbed for kinematics evaluation.

For the ease of calculation, the adjacent rotating axis of each joint is perpendicular to each other. Hence, the link length is also the length of the link body. Beside, those rotating axis will always have link twist of 90 degree.

The configuration comprises 6 frames (5 for joints and one for the end-effector) and 5 links in total. This structure is flexible to do complicated tasks, while remains simple to study its kinematics.

### 2.2 Denavit-Hartenberg (DH) parameters

Frame transformations are calculated by applying translating and rotating matrices. DH parameters are used in finding the kinematics solutions of our arm. Below is the DH parameter description of the arm model:

where *q*_*n*_ is the position angle of the servo at that joint.

### 2.3 Constraints

There are orientation constraints of the coordinate systems:

- The *z* axis is in the direction of the joint axis. The servo motors will rotate about this axis.
- The adjacent joint x axis is parallel to the common normal of the current z axis and the previous joint *z* axis: *x*_*n*_ = *z*_*n*_ *× z*_*n*−1_. This means it is perpendicular to both *z* axes
- The *y* axis completes a right-hand reference frame based on *x* and *z*

**TABLE 1.** DH parameters

| Joint | $d_n$ | $\theta_n$ | $r_n$ | $\alpha_n$ |
| --- | --- | --- | --- | --- |
| 0 | $h$ | $q_0 + \pi$ | 0 | $\frac{\pi}{2}$ |
| 1 | 0 | $q_1 + \frac{\pi}{2}$ | $l_1$ | $\frac{\pi}{2}$ |
| 2 | $h$ | $q_2$ | 0 | $-\frac{\pi}{2}$ |
| 3 | 0 | $q_3 + \frac{\pi}{2}$ | $l_2$ | $\frac{\pi}{2}$ |
| 4 | $l_3$ | $q_4$ | 0 | 0 |

## 3 Adaptive Neural Fuzzy Inference System (ANFIS) approach of Inverse Kinematics

Beside numerical method that calculates joint angles directly from 3D pose of end-effector in straight-forward manner, in this section, we present ANFIS approach that has learning and highly adaptation capability depending on workspace conditions. This technique, which bases on soft-computing, is a better choice for real-time control application due to efficient computing characteristic. There are some previous applications of ANFIS applied for inverse kinematics problems, for instances, ANFIS approach has been successfully applied for 3DOF planar manipulator in [9] with under 10% error and 3DOF SCARA manipulator(two revolute joints and a prismatic joint) [10]. For our arm model, we build an appropriate ANFIS model that applies for 4DOF human-like arm in 3-dimension spherical coordinates. The goal of this approach is to map Cartesian coordinate in to joint angle vector space, which learns from dataset produced by forward kinematics.

### 3.1 ANFIS

ANFIS combines the strength of two soft computing frameworks of Artificial Neural Network (ANN) and Fuzzy Logic [11], in which the power of Fuzzy Logic is parameterizing categories for process of quantitative analysis and ANN is the learning ability as a guide for adjusting membership functions (MFs) of Fuzzy Logic. ANFIS has higher learning ability than ANN and therefore produces smaller error in inverse kinematics. The detail architecture of ANFIS is described in Figure 4.

**FIGURE 1.**
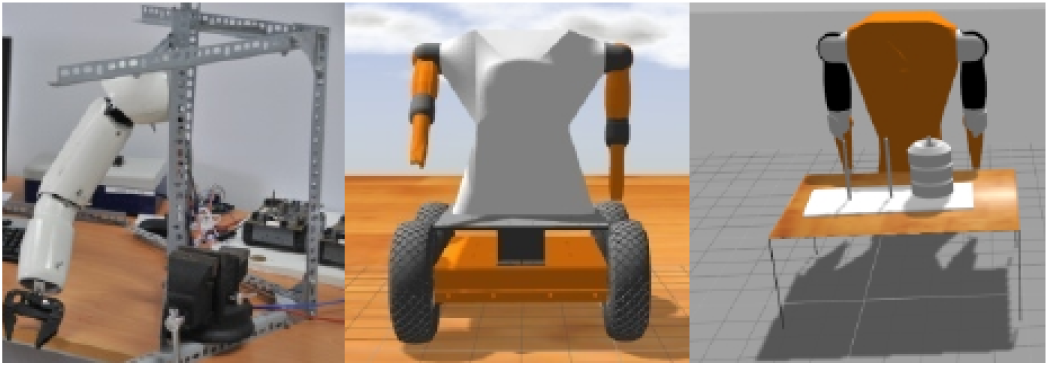
Our Simulator for Mobile robot manipulators

**FIGURE 2.**
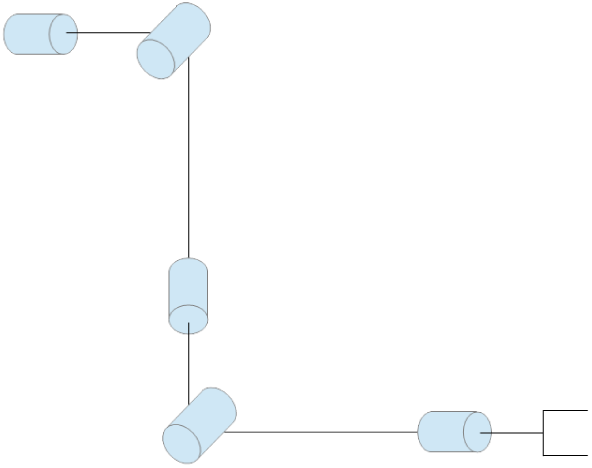
Simple model of series rotation axes

**FIGURE 3.**
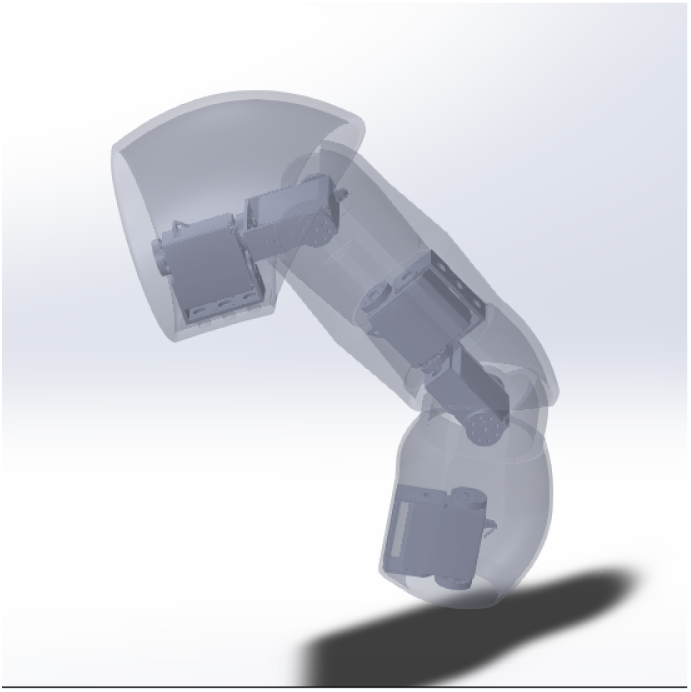
Arm design with servo motor positions

**FIGURE 4.**
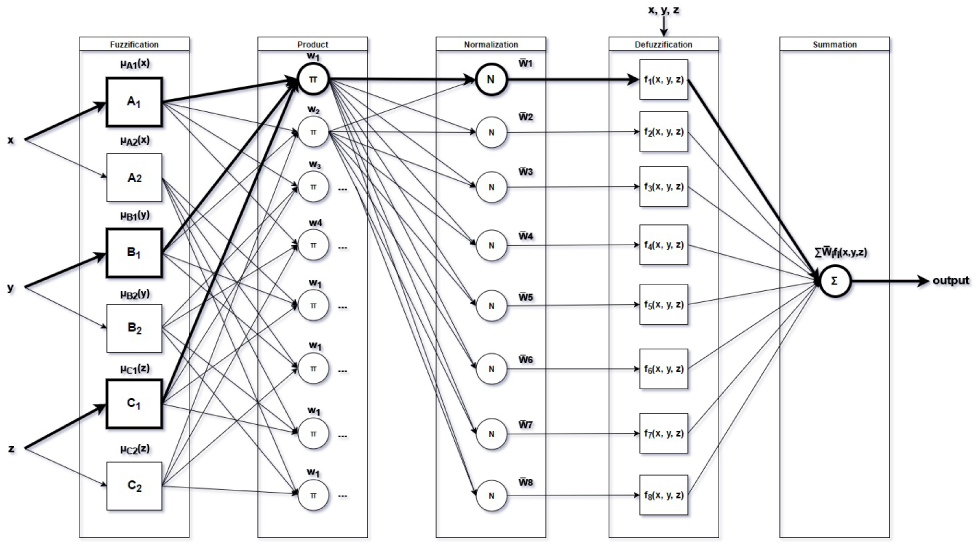
ANFIS architecture of 3 input parameter and 8 rules

In general, the architecture of type-3 ANFIS is an adaptive network that uses feed-forward neural network to learn premise and consequent parameters of Fuzzy Logic framework. Type-3 ANFIS adapts Takagi-Sugeno if-then rules as the following 8 rules in Figure 4:

- **Rule 1: IF x is A1, y is B1 and z is C1 THEN** *f*_1_ = *p*_1_*x* + *q*_1_*y* + *t*_1_*z* + *r*_1_
- Rule 2: IF x is A1, y is B1 and z is C2 THEN *f*_2_ = *p*_2_*x* + *q*_2_*y* + *t*_2_*z* + *r*_2_
- Rule 3: IF x is A1, y is B2 and z is C1 THEN *f*_3_ = *p*_3_*x* + *q*_3_*y* + *t*_3_*z* + *r*_2_
- …
- Rule 8: IF x is A2, y is B2 and z is C2 THEN *f*_8_ = *p*_8_*x* + *q*_8_*y* + *t*_8_*z* + *r*_8_

The list of rules is generated by the combination of member functions corresponding to each input. For simplicity, we put et cetera in rule list and architecture diagram due to pattern. Note that the highlighted **Rule 1** corresponds to the highlighted edges in Figure 4 emphasizing Rule 1.

ANFIS has five layers, the first and the fourth are adaptive layers, others contain fixed nodes. A description of these layers is described as following:

- **Fuzzification layer**: Each node in this layer contains membership function

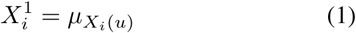

with linguistic label *X*^1^ = (*A, B, C*), *i* is MF number each input parameter *u* = (*x, y, z*). In the case of our system, we choose 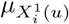 as Gaussian function due to the characteristic of Cartesian space and faster training:

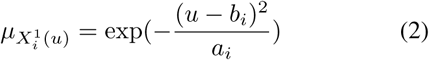

where *a*_*i*_, *b*_*i*_ are premise parameters for this layer.
- **Product Layer**: Nodes in this layer are circle labeled *π*, which means they are fixed. These nodes are T-norm operator that performs AND characteristic. For simplicity, these nodes just multiplies incoming signals and output firing strength:

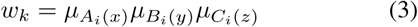
- **Normalization layer**: Each node in this layer are also fixed node labeled *N*. *kth* node normalizes *kth* firing strength to the sum of firing strength:

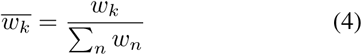
- **Defuzzification layer**: Each node in this layer is a THEN term of Takagi-Sugeno rule, they receive inputs *x, y, z* and normalized firring strength and send out linear combination of inputs:

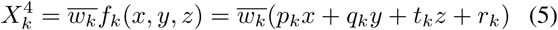

where (*p*_*k*_, *q*_*k*_, *t*_*k*_, *r*_*k*_) are consequent parameters that will be trained by supervised learning algorithm.
- **Summation layer**: This layer contains single node that sums all incoming signals from previous layer:

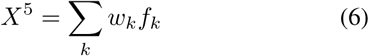

### 3.2 Training Applied Inverse Kinematics Framework

Due to the property of ANFIS that predicts only one output from set of inputs, we constructs the inverse kinematic framework with four ANFIS components corresponding to four joint angles of robotic arm. Each ANFIS is inputed with 3-dimension Cartesian coordinates (*x, y, z*) and outputs mapped joint angles (*q*_1_, *q*_2_, *q*_3_, *q*_4_) (Figure 5).

**FIGURE 5.**
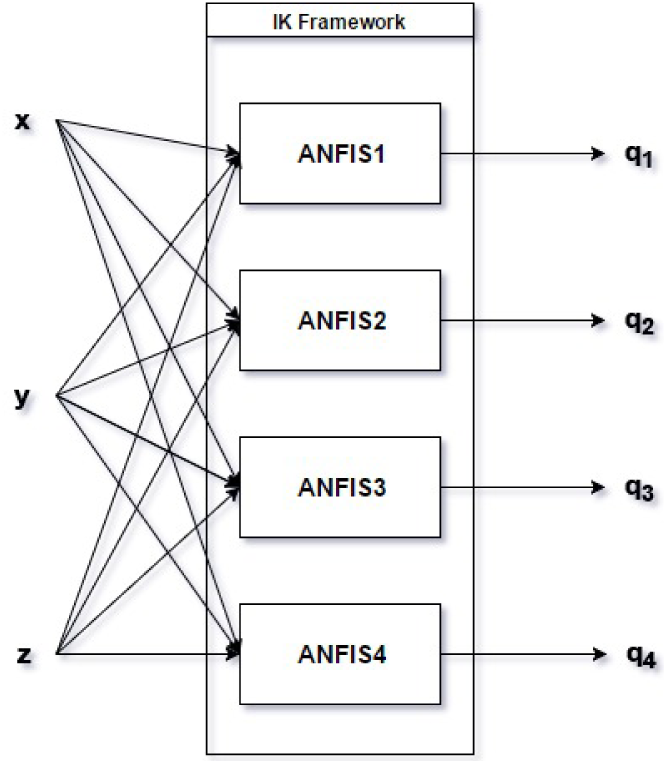
Applied Inverse Kinematic Framework for Human-like robotic arm

**FIGURE 6.**
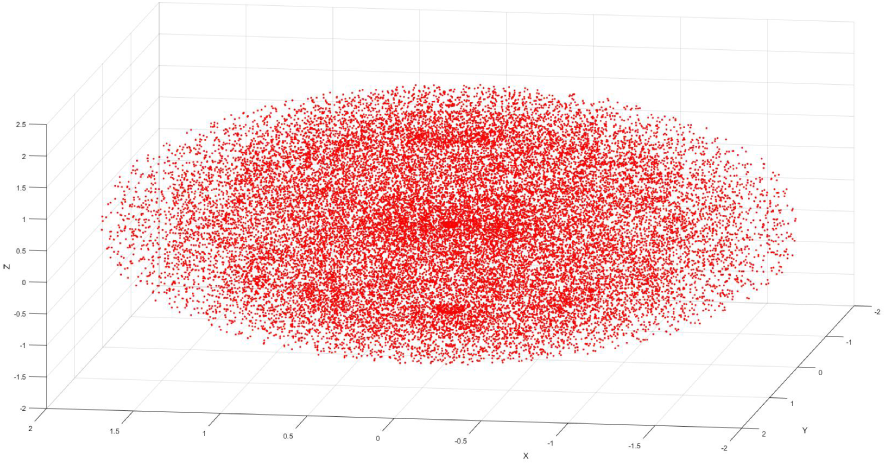
Workspace of end-effector with no constraint in joint angles

A training set that maps 3-dimension coordinates to joint-angles is produced by forward kinematic. the set is then split to four training subsets, each subset consists of coordinates of end-effector and a joint angle. These subsets are used for training each ANFIS component corresponding to each joint angle that will result end-effector pose.

In ANFIS architecture, Fuzzification and Defuzzification step contain modifiable nodes that are the training targets. We can use backpropagation algorithm to train the parameters of both layers. However, backpropagation algorithm has been found problematic that tends to converge to local minima and has a slow convergence rate. A hybrid training method is propose by [**?**], the method uses combination of backpropagation algorithm to train non-linear premise parameters of the first layer and Recursive Least Square Estimator (RLSE) to train linear consequent parameter of Defuzzification layer. one iterative of the method consists of forward path and backward path.

In forward path, premise parameters is steady, RLSE is applied for consequent parameters to gradually repair their values. After the consequent parameters are fixed, the inputs data is applied to the framework in order to generate the output that are then compared to desired output to calculate error.

In backward path, the consequent parameters are steady, the errors are propagated back to repair the premise parameters by using backpropagation algorithm.

The combination of backpropagation and RLSE produces smaller errors to the desired output and faster convergence comparing to single backpropagation algorithm for learning two kind of parameters. The detail of hybrid method can be refer in (cite Jang).

## 4 Model Evaluation

This section will provide the evaluation of our arm model as well as its Inverse Kinematic (IK) solutions. Our models will be evaluated its coverage as workspace, our IK solution error using IK solution using ANFIS.

### 4.1 Workspace

At first, we setup our test environment on MATLAB with the configuration of the arm model as DH parameters below:

**TABLE 2.** Experiment DH parameters

| Joint | $d_n$ | $\theta_n$ | $r_n$ | $\alpha_n$ |
| --- | --- | --- | --- | --- |
| 0 | 0.1 | $q_0$ | 0 | $\frac{\pi}{2}$ |
| 1 | 0 | $q_1$ | 1 | $\frac{\pi}{2}$ |
| 2 | 0.1 | $q_2$ | 0 | $-\frac{\pi}{2}$ |
| 3 | 0 | $q_3$ | 1 | $\frac{\pi}{2}$ |

To clarify, *d*_*n*_ is the small gap between two joint servo motors, *r*_*n*_ corresponds to the segment lengths of the arm between links.

To visualize the workspace, a 4-dimension grid of four joint angles (*q*_0_, *q*_1_, *q*_2_, *q*_4_) is generated with no constraint *q*_*n*_ = 0 : 2*π*, step 0.3 radian. The grid is then iterated over its joint angle values, these values are passed to forward kinematics solution to project all coordinates that the end-effector can reach. The workspace of the arm is shown in Figure as red cloud points below:

The shape of workspace is ellipsoid with volume *V* = 29.59 calculated from DH parameters, which means our model can reach more coordinates in horizontal direction than vertical direction. The model demonstrates its feasibility due to most environmental workspaces in reality are also ellipsoid.

In the case of real arm, the constraints are applied to each joint angles of each link. For the case of servo motors, the constraint 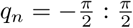 is applied for all joint angles with the same angle resolution. Figure 7 describes workspace in this case.

**FIGURE 7.**
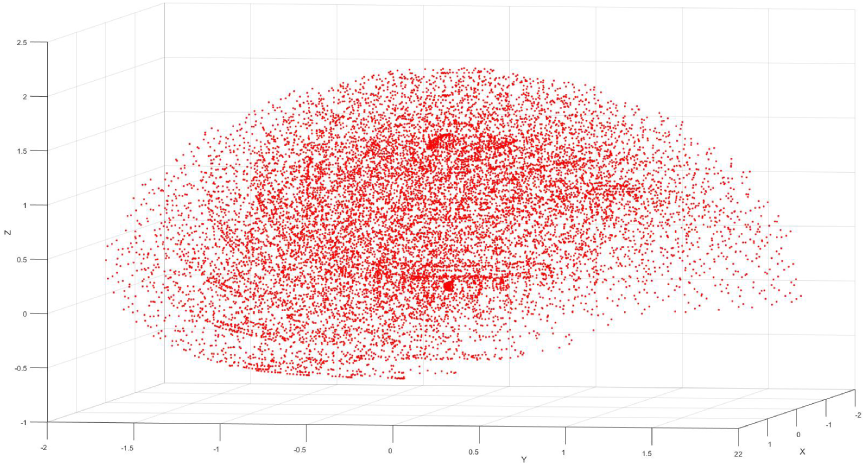
Workspace of end-effector with 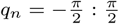 constraint in joint angles

**FIGURE 8.**
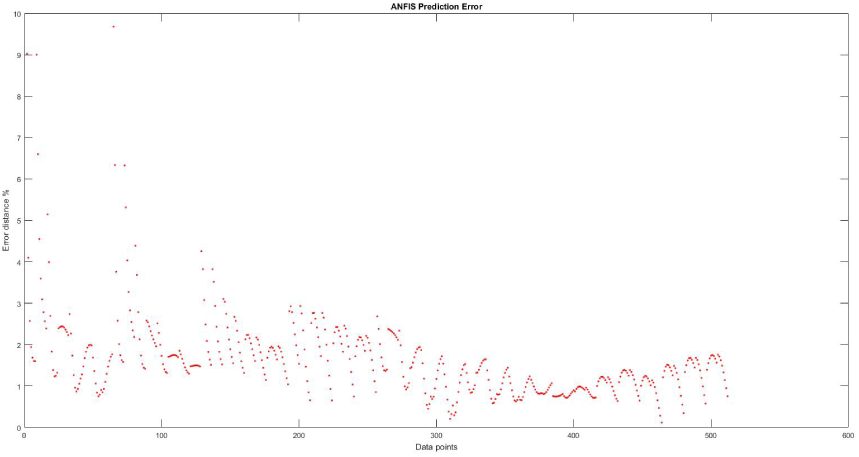
Error evaluation with training parameter MFs = 3 and mean error percentage = 1.94%

**FIGURE 9.**
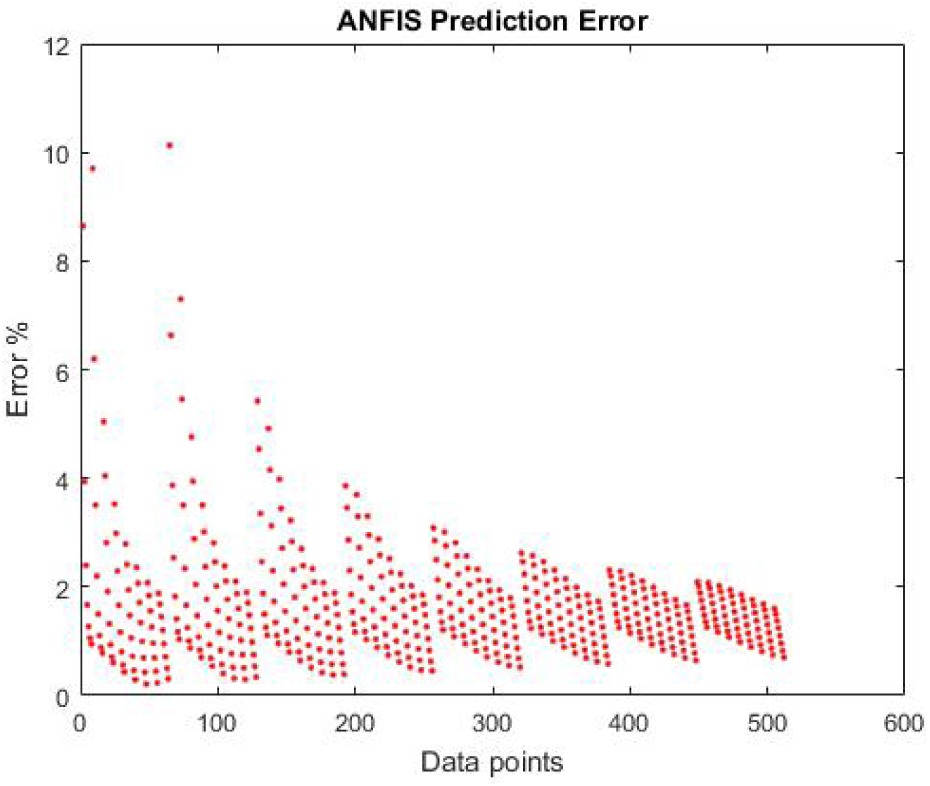
Error evaluation with training parameter MFs = 5 with mean error percentage = 1.65%

The figure now has snail-shell shape with volume *V* = 13.80. Total volume of accessible space is reduced by half due to joint angle space volume is 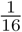 the no-constraint volume.

The next section will demonstrate the solution error of each mentioned IK method.

### 4.2 Error Evaluation of ANFIS approach

In this approach, we generate 65536 data points as 4D grid joint angles space described as training set for four ANFIS components. We establish evaluation metric by generating 3-dimension coordinate grid (*x, y, z*)

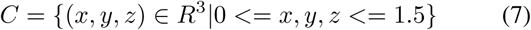

as 1.5*x*1.5*x*1.5 cubic space, step 0.2 unit length, within model workspace. The coordinate grid consists of 512 data points and is then used to calculate joint angle space by each IK methods. After that, the result joint angle space is passed to forward kinematic solution to project back result Cartesian coordinate, initial coordinates and result coordinates will be used to calculate IK coordinate error percentage of each method. This evaluation metric ensures generalization of ANFIS evaluation due to different training set and evaluation set. We evaluate the performance of this IK approach by training ANFIS with increasing member functions and epochs size, then apply evaluation metric described.

As can be seen, both evaluation results show that ANFIS has reliable and accurate IK solutions with error lower than 2% and average computed time only 0.5ms each data point in term of performance. The first evaluation has 0.026 mean distance value and 0.007 standard deviation. In second case, mean distance error is 0.022 and standard deviation is 0.006. Prediction data in second evaluation is more steadily due to increased member function and epoch size. There are spikes up to 12% in both cases due to the boundary conditions of test set. Training set with higher resolution than test set can resolve the problem.

## 5 Conclusion

This document proposes robust and inexpensive robotic arm model with detail description. The model can be applied with different kind of motors and arm segment material.

Experiments also shows applied learning methods, particularly ANFIS, has robust application in solving inverse kinematic problem. Despite of 47 hours training time with over 65536 data points, the proposed applied ANFIS produces result rapidly with low error rate below %2. This framework also implies ANFIS can be applied for more DOFs of arm in 3-dimension space with orientation constraints.

In our future work, we will study applying our approaches using vector Calculus and ANFIS framework with higher DOF arm model with orientation parameter and wider joint angle space. There are the other works need to improve mechanism such as variable stiffness joints while it contains the limitations of motor actuators.

## References

[1] H. V. Nguyen, T. D. Le, D. D. Huynh, P. Nauth, Forward kinematics of a human-arm system and inverse kinematics using vector calculus, in: The 14th International Conference on Control, Automation, Robotics and Vision (ICARCV) 2016, 2016, pp. 16. doi: 10.1109/ICARCV.2016.7838641.

[2] G. Antonio, M. Manuel, Vector Analysis Versus Vector Calculus, Springer Publishing Company, Incorporated, 2012. doi: 10.1007/978-1-4614-2200-6.

[3] D. Shin, I. Sardellitti, Y.-L. Park, O. Khatib, M. R. Cutkosky, Design and control of a bio-inspired human-friendly robot 29 (5) (2010) 571584.

[4] A. Miller, P. Allen, V. Santos, F. Valero-cuevas, From robot hands to human hands: A visualization and simulation engine for grasping research, Industrial Robot 32 (2005) 5563.

[5] M. Grebenstein, A. Albu-Schffer, T. Bahls, M. Chalon, O. Eiberger, W. Friedl, R. Gruber, S. Haddadin, U. Hagn, R. Haslinger, H. Hppner, S. Jrg, M. Nickl, A. Nothhelfer, F. Petit, J. Reill, N. Seitz, T. Wimbck, S. Wolf, T. Wsthoff, G. Hirzinger, The dlr hand arm system, in: 2011 IEEE International Conference on Robotics and Automation, 2011, pp. 31753182. doi: 10.1109/ICRA.2011.5980371.

[6] I. Dulba, M. Opaka, A comparison of jacobian-based methods of inverse kinematics for serial robot manipulators, in: International Journal of Applied Mathematics and Computer Science, 2013. doi: 10.2478/amcs-2013-0028.

[7] E. Oyama, N. Y. Chong, A. Agah, T. Maeda, Inverse kinematics learning by modular architecture neural networks with performance prediction networks, in: Proceedings 2001 ICRA. IEEE International Conference on Robotics and Automation (Cat. No.01CH37164), Vol. 1, 2001, pp. 1006 1012 vol.1. doi: 10.1109/ROBOT.2001.932681.

[8] S. Tejomurtula, S. Kak, Inverse kinematics in robotics using neural networks, Information Sciences 116 (2) (1999) 147 164. doi: http://dx.doi.org/10.1016/S0020-0255(98)10098-1.

[9] A.-V. Duka, Anfis based solution to the inverse kinematics of a 3dof planar manipulator, 533, Procedia Technology 19 (2015) 526 8th International Conference Interdisciplinarity in Engineering, INTER-ENG 2014, 9-10 October 2014, Tirgu Mures, Romania. doi: http://dx.doi.org/10.1016/j.protcy.2015.02.075. URL http://www.sciencedirect.com/science/article/pii/S2212017315000766

[10] W. M. Jasim, Solution of inverse kinematics for scara manipulator using adaptive neuro-fuzzy network, 2011.

[11] J. S. R. Jang, Anfis: adaptive-network-based fuzzy inference system, IEEE Transactions on Systems, Man, and Cybernetics 23 (3) (1993) 665685. doi: 10.1109/21.256541.

